# Organelle-Aware Representation Learning Enables Label-Free Detection of Mitochondrial Dysfunction in Live Human Neurons

**DOI:** 10.1101/2025.10.12.681848

**Authors:** SeungJu Yoo, Eunjin Yang, Seo-Hyun Kim, Hoewon Park, Daesoo Kim, Ki-Jun Yoon, Minee L. Choi

## Abstract

Mitochondrial dysfunction is a convergent hallmark of neurodegenerative diseases and represents a promising biomarker for early diagnosis and therapy. However, current in vitro assays rely on fluorescence or electron microscopy, which are invasive, low-throughput, and incompatible with longitudinal analysis. Here, we present a noninvasive framework by integrating label-free optical diffraction tomography (ODT) with organelle-aware representation learning to detect subtle mitochondrial dysfunction in live human induced pluripotent stem cell (hiPSC)-derived neurons. Through virtual staining of the nuclei, lysosomes, and mitochondria, we establish two complementary and interpretable classification pipelines: a deep learning model with organelle-aware encoder and a logistic regression model on morphometric descriptors. Both models achieve approximately 85% accuracy: the deep model provides end-to-end prediction with minimal feature engineering, whereas the logistic regression model offers a more interpretable, feature-based approach. To our knowledge, this is the first demonstration of ODT-based organelle-resolved virtual staining in live human neurons, establishing a scalable, non-invasive platform for mitochondrial disease modeling, drug discovery, and neurodegeneration research.

## Introduction

Neurodegenerative diseases, including Parkinson’s disease and Alzheimer’s disease are characterized by progressive and irreversible loss of neuronal function, representing a growing global health burden^1,2^. Although each disorder arises from distinct and multifactorial mechanisms, mitochondrial dysfunction is a convergent hallmark, manifesting as impaired oxidative phosphorylation, elevated oxidative stress, and disrupted cellular homeostasis^3–8^. Importantly, mitochondrial abnormalities often precede overt neuronal loss, positioning them as promising early biomarkers and therapeutic targets^9–12^.

Human induced pluripotent stem cell (hiPSC)-derived neurons provide a powerful system for modeling patient-specific disease phenotypes and evaluating candidate therapies while retaining the genetic background associated with disease susceptibility^13–18^. These platforms have successfully recapitulated mitochondrial dysfunction observed in patients, creating opportunities for mechanistic insights and drug discovery. However, current assays for mitochondrial health in iPSC-derived neurons typically rely on fluorescent dyes, genetically encoded reporters, or electron microscopy. Such methods are invasive, perturb physiology, and are incompatible with high-throughput or longitudinal studies^19–21^. This limitation underscores the need for non-invasive, label-free approaches capable of sensitively detecting mitochondrial dysfunction in live human neurons.

Optical diffraction tomography (ODT), also known as holotomography, is a label-free, three-dimensional imaging modality that leverages refractive index (RI) contrast to reconstruct quantitative phase tomograms of living cells^22–25^. Unlike fluorescence microscopy, ODT provides intrinsic biophysical information without exogenous labels, avoiding phototoxicity and allowing repeated imaging over time. Previous studies have demonstrated the utility of ODT in structural analysis, monitoring cell death, and dynamic organelle behavior across various cell types including fibroblast, white blood cells, neurons, and cancer cells^26–32^. However, its application to hiPSC-derived neurons particularly in the context of disease modeling and prediction remains largely unexplored due to their fine neurites and compact somata.

We present a label-free, deep-learning framework that extracts biologically meaningful features from ODT volumes of live hiPSC-derived neurons. A multi-label virtual-staining network segments nuclei, mitochondria, and lysosomes, yielding organelle-aware representations that drive two complementary, interpretable classifiers: an end-to-end model built on the organelle-aware encoder and a descriptor-based logistic regression using morphometrics from predicted masks (Fig. 1A). Both approaches sensitively detect subtle mitochondrial dysfunction induced by electron transport chain (ETC) inhibitors, and their concordant predictions provide mutual cross-validation and highlight that organelle-representations drive the detection. Together, these pipelines enable non-invasive, high-throughput detection of mitochondrial dysfunction with transparent mechanistic readouts. To our knowledge, this is the first organelle-resolved virtual staining of hiPSC-derived neurons with ODT, establishing a scalable platform for longitudinal monitoring of mitochondrial health, therapeutic screening, and mechanistic studies of neurodegeneration.

**Figure 1.**
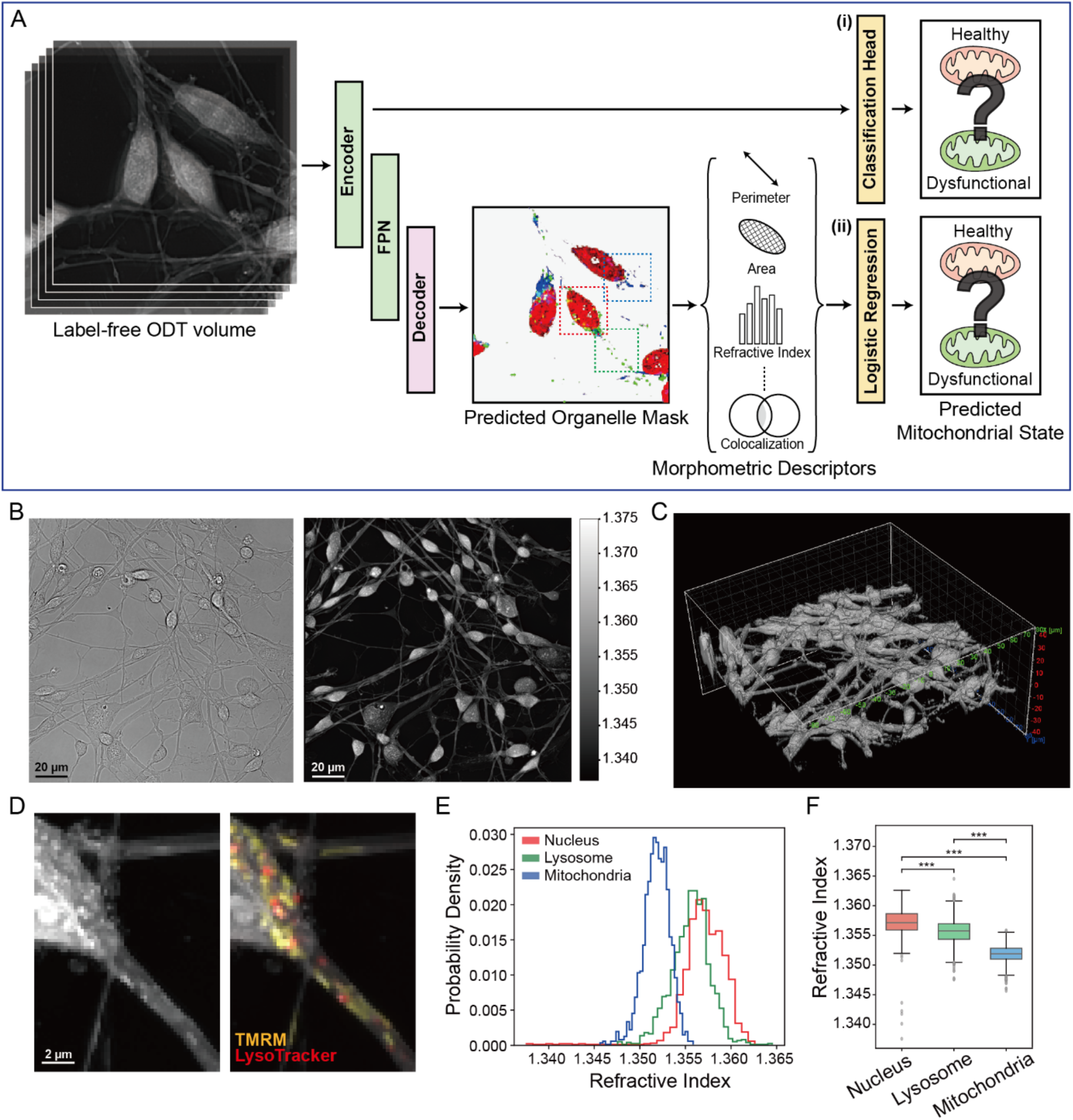
Label-free imaging using optical diffraction tomography (ODT). (A) A schematic overview. Virtual staining of subcellular organelles in hiPSC-derived neurons from ODT. Mitochondrial dysfunctions can be identified through two independent and complementary classification approaches. (B) Direct comparison of standard brightfield image and ODT (2D projection). Subcellular features in transparent cells cannot be resolved from the brightfield image. (C) Reconstructed 3D volume of the cells shown in (B). (D) Correlative fluorescent imaging showing mitochondria (TMRM) and lysosomes (LysoTracker) as distinguishable features in the refractive-index (RI) map. (E) The probability density of pixel RI of subcellular organelles extracted from ODT using fluorescent images. (F) The distribution of pixel RI of each organelle analyzed from 1480 ODT volumes. Statistical analysis performed using Student’s t-test with Holm-Bonferroni correction (***: p < 0.001).

## Results

### Optical Diffraction Tomography Resolves Subcellular Structure in Live Neurons

Conventional brightfield microscopy, while widely accessible, provides only superficial visualization of cellular morphology. Its utility is limited by the inherent transparency of biological cells and tissues, which obscures subcellular structures. By contrast, label-free refractive index (RI)-based optical diffraction tomography (ODT) enables high-contrast visualization of transparent cells (Fig. 1B-C). While single RI value does not map to a specific subcellular structure, organelles such as mitochondria can be distinguished qualitatively based on morphology and spatial context (Fig. 1D). Analysis of RI distributions of nuclei, lysosomes, and mitochondria via fluorescence alignment found that each organelle, especially mitochondria, occupied characteristic range in the RI spectrum (Fig. 1E-F). This observation motivates subsequent claims that ODT volumes contain sufficient subcellular information, and it can be readily utilized through organelle-aware representation learning.

### Generation of Mitochondrial Dysfunction Model from hiPSC-Derived Cortical Neurons

Human neurons differentiated in vitro are particularly sensitive to variability in cell state, making it essential to establish a robust baseline of healthy, mature cortical neurons prior to modeling mitochondrial dysfunction. Cortical neurons were generated from hiPSCs via dual SMAD inhibition (Fig. 2A)^33,34^ and maintained until day 63 (D63). At this stage, cellular identity and functional maturity were confirmed. Immunocytochemistry revealed strong expression of neuronal markers MAP2 and TuJ1, as well as the cortical upper-layer marker SATB2, consistent with successful differentiation into mature cortical neurons (Supplementary Fig.1A-B). Their functional competence was validated by calcium imaging using the intracellular calcium indicator, Fluo-4. Depolarization with potassium chloride (KCl; 50μM) induced a rapid increase in fluorescence intensity, demonstrating the presence of functional voltage-gated calcium channels (Fig. 2B-C). Mitochondrial metabolic activity was further assessed by live-cell imaging with tetramethylrhodamine (TMRM). Cells exhibited strong baseline fluorescence reflecting intact membrane potential, which dissipated rapidly following treatment with the mitochondrial uncoupler FCCP (Fig. 2D-E). Together, these results confirm that the differentiated neurons possessed appropriate marker expression, physiological activity, and intact mitochondrial function, providing a robust foundation for controlled perturbation experiments.

**Figure 2.**
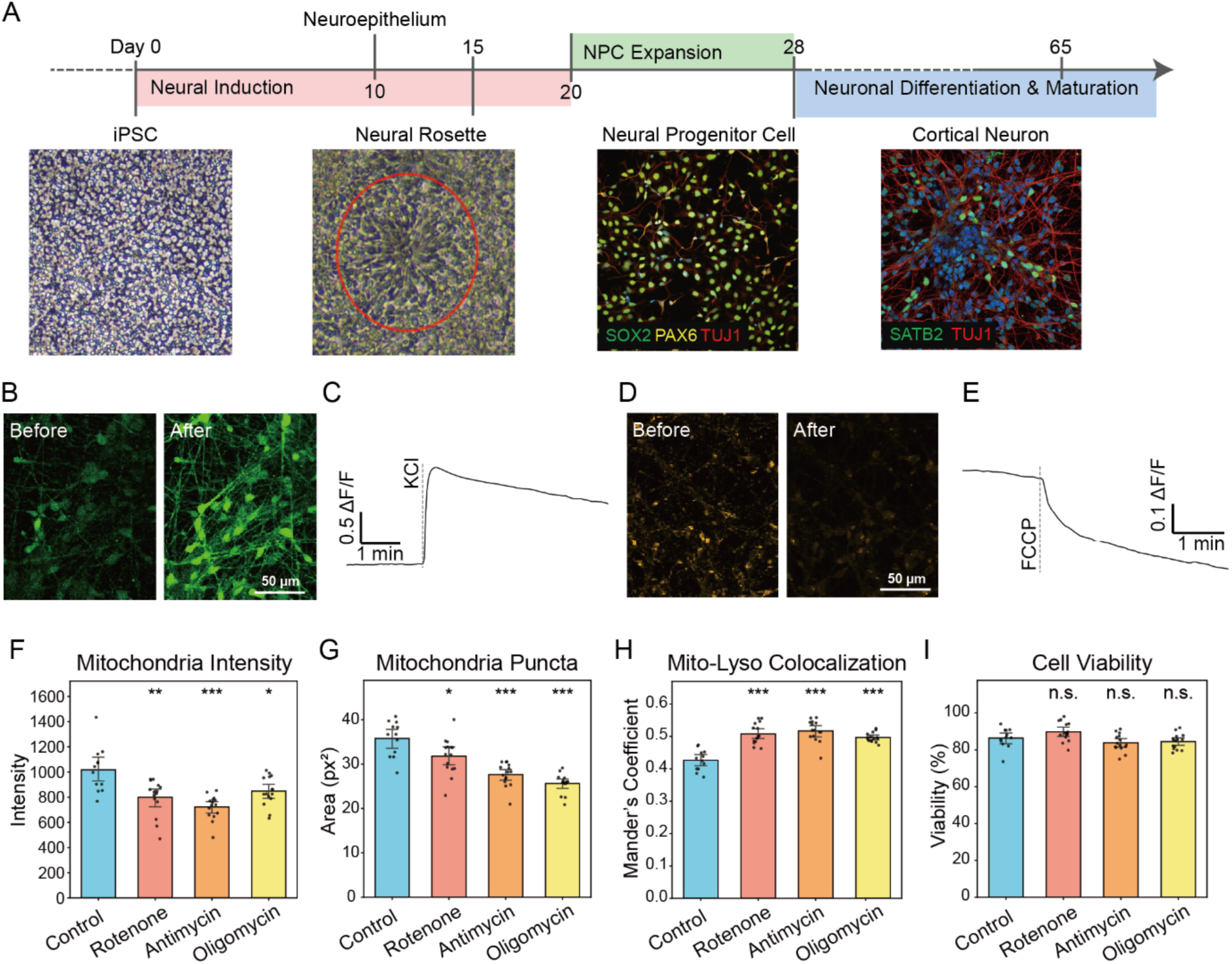
Generation of a mitochondrial dysfunction model by ETC inhibition on hiPSC-derived neurons. (A) Timeline of neuronal differentiation. The protocol follows 10 days of dual-SMAD inhibition and the generation of a neuroepithelial sheet, followed by the neural rosettes at around day 15. Neurons are cultured for at least 65 days post-induction to ensure maturity and functionality. Representative images are shown for each stage. (B-C) Calcium imaging using the intracellular calcium indicator, Fluo-4. A sharp increase in fluorescence intensity after KCl treatment reflects membrane depolarization leading to calcium influx through voltage-gated calcium channels. (D-E) Strong baseline TMRM intensity representing functional mitochondria maintaining mitochondrial membrane potential. Mitochondrial uncoupler FCCP treatment quickly dissipates the potential. (F-I) Live cell analysis post 24 hours of mitochondrial dysfunction inducers. (F) Lower mitochondrial intensity denotes reduced membrane potential, (G) fewer and smaller puncta indicate fragmentation, (H) increased mitochondria–lysosome overlap suggests enhanced mitophagy, and (I) cell viability remained unaffected after chemical treatment, confirming preserved overall cellular function.

To model mitochondrial dysfunction, we applied established electron transport chain (ETC) inhibitors (rotenone, antimycin, and oligomycin) which selectively impair mitochondrial bioenergetics and increase oxidative stress^35–42^. The use of defined chemical inhibitors provides a standardized and precise means of inducing mitochondrial dysfunction, in contrast to patient-derived cells where multiple pathological features coexist and complicate the attribution of specific morphological changes. Such biological complexity in patient-derived models introduces label noise at training phase, potentially obscuring consistent morphological cues required for deep learning approaches.

The effects of mitochondrial perturbation were validated by fluorescent imaging of mitochondria and lysosomes using TMRM and LysoTracker, respectively. Treatment reduced mitochondrial membrane potential, as reflected in decreased TMRM intensity (Fig. 2F), and promoted mitochondrial fragmentation, evidenced by reduced puncta size (Fig. 2G). In addition, colocalization analysis demonstrated an increase in the TMRM–LysoTracker overlap (Fig. 2H), consistent with enhanced mitophagy and clearance of damaged mitochondria.

Importantly, we used low concentrations at 100 nM of ETC inhibitors and limited exposure to 24 hours to preserve overall neuronal viability (Fig. 2I). This approach models the subtle mitochondrial dysfunction characteristic of early neurodegenerative disease stages, while minimizing nonspecific cellular deterioration. Such controlled perturbation is also critical for downstream neural network training, as severe functional collapse or cell death could bias models toward learning spurious features of cell stress rather than mitochondrial-specific dysfunction.

### Organelle-Aware Representation Learning via Virtual Staining

Detecting mitochondrial dysfunction with ODT rests on two premises: (a) ODT provides sufficient information to delineate organelles, and (b) organelle morphology and context are predictive of mitochondrial state. The correlation between mitochondrial structure and function is well established^43–46^. However, whether optical diffraction tomography (ODT) contains sufficient subcellular detail in live neurons remained uncertain.

To further investigate the latent information in ODT, we designed a virtual staining model that can extract subcellular information to resolve three key organelles (nuclei, lysosomes, mitochondria) concerning mitochondrial dysfunction. We adopt a 2.5D, multi-label segmentation architecture based on a ResNet-34 encoder^47^ and FPN-inspired^48^ multi-scale aggregator to produce per-organelle segmentation predictions (Fig. 3A). A total of 1,622 ODT volumes were pooled across multiple experimental states (see Methods section for segmentation data composition), and each volume consisted of 88 optical sections (741 × 741 pixels each) with voxel dimensions of 1 × 0.222 × 0.222 μm³. We designed the input dimension as 7 × 224 × 224 pixels, corresponding to seven 50 × 50 μm cross-sections, each containing approximately 4-6 neurons. This balances the computational overhead of processing the full volume while retaining sufficient structural context of neuronal networks. Ground truth annotations were generated using correlated fluorescent imaging of each organelle and reduced to 2D by maximum intensity projection (MIP) to mitigate the spatial misalignment between the two imaging modalities across depth-axis The trained model produced reliable segmentation masks for the three organelles (Fig. 3B & Supplementary Fig. 2A), yielding Dice coefficients of 0.92 (nuclei), 0.43 (lysosomes), and 0.48 (mitochondria) (Fig. 3C). While Dice is widely used for evaluating segmentation performance, it is sensitive to class imbalance and uncertainty in ground truth masks. Lysosome and mitochondria pixels comprise <5% of the ODT image and temporal misalignment between ODT and fluorescence caused by organelle motility during acquisition contributed to low Dice scores. By adopting peak-based instance matching, the lysosome and mitochondria showed high precision and low recall suggesting the segmented pixels are highly reliable (few false positives), but less confident regions of true positive pixels are missed (Supplementary Fig. 2B). Additionally, relaxed Dice was computed after dilating prediction and ground-truth masks by 3 pixels (666 nm), increasing the metrics to 0.53 for lysosomes and 0.65 for mitochondria (Fig. 3C). Qualitatively, predictions localized organelles as probabilistic regions rather than pixel-accurate boundaries, consistent with uncertainty in the ground truth due to motility and spatial variability.

**Figure 3.**
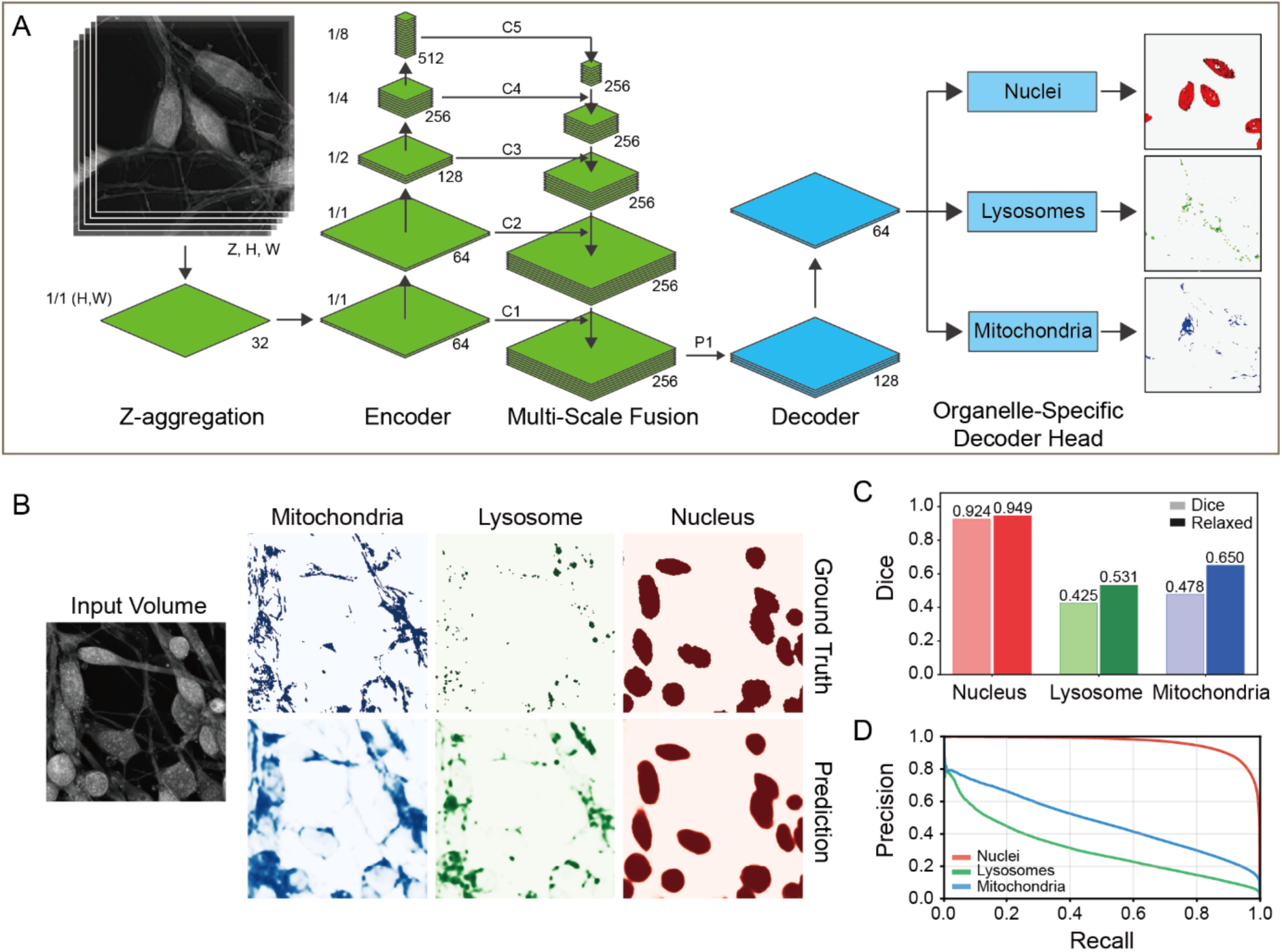
Segmentation of subcellular organelles in hiPSC-derived neurons from ODT. (A) Architecture of the segmentation model used for this study. ODT volume is preprocessed by a 2.5D convolution block to collapse the 3D volume. Encoder follows ResNet34-based architecture with FPN-based multi-scale fusion layers. Segmentation masks are generated from each organelle-specific decoder head. (B) Representative segmentation images. The input 3D volume is shown as a 2D maximum intensity projection. (C) Segmentation performance on the test dataset. Mitochondria and lysosomes segmentation performance is better represented by the relaxed Dice score, which dilates the ground truth (GT) and the prediction mask by 3 pixels (222 nm). This accounts for the motility of the organelles and temporal misalignment between ODT and fluorescent ground truths. (D) Precision-recall curve for each organelle segmentation.

These results demonstrate that ODT not only encodes structural information at the subcellular level but also enables localization of the organelles. ODT thus provides a viable, label-free modality for monitoring organellar morphology, and establishes a methodological foundation for subsequent classification of mitochondrial dysfunction in live neuronal systems without reliance on exogenous labels.

### Organelle-Aware Encoder Enables Classification of Mitochondrial Dysfunction

Using the chemically induced human neuronal model of mitochondrial dysfunction, we acquired 1020 ODT volumes and trained a binary classification head attached directly to the organelle-aware encoder trained from organelle segmentation. The head consisted of a multi-head attention pooling module followed by a multilayer perceptron, operating on the highest-level C5 features of the ResNet-FPN backbone (Fig. 4A). During training, all segmentation weights were frozen, ensuring that classification was based exclusively on organelle-aware features while preserving segmentation fidelity.

**Figure 4.**
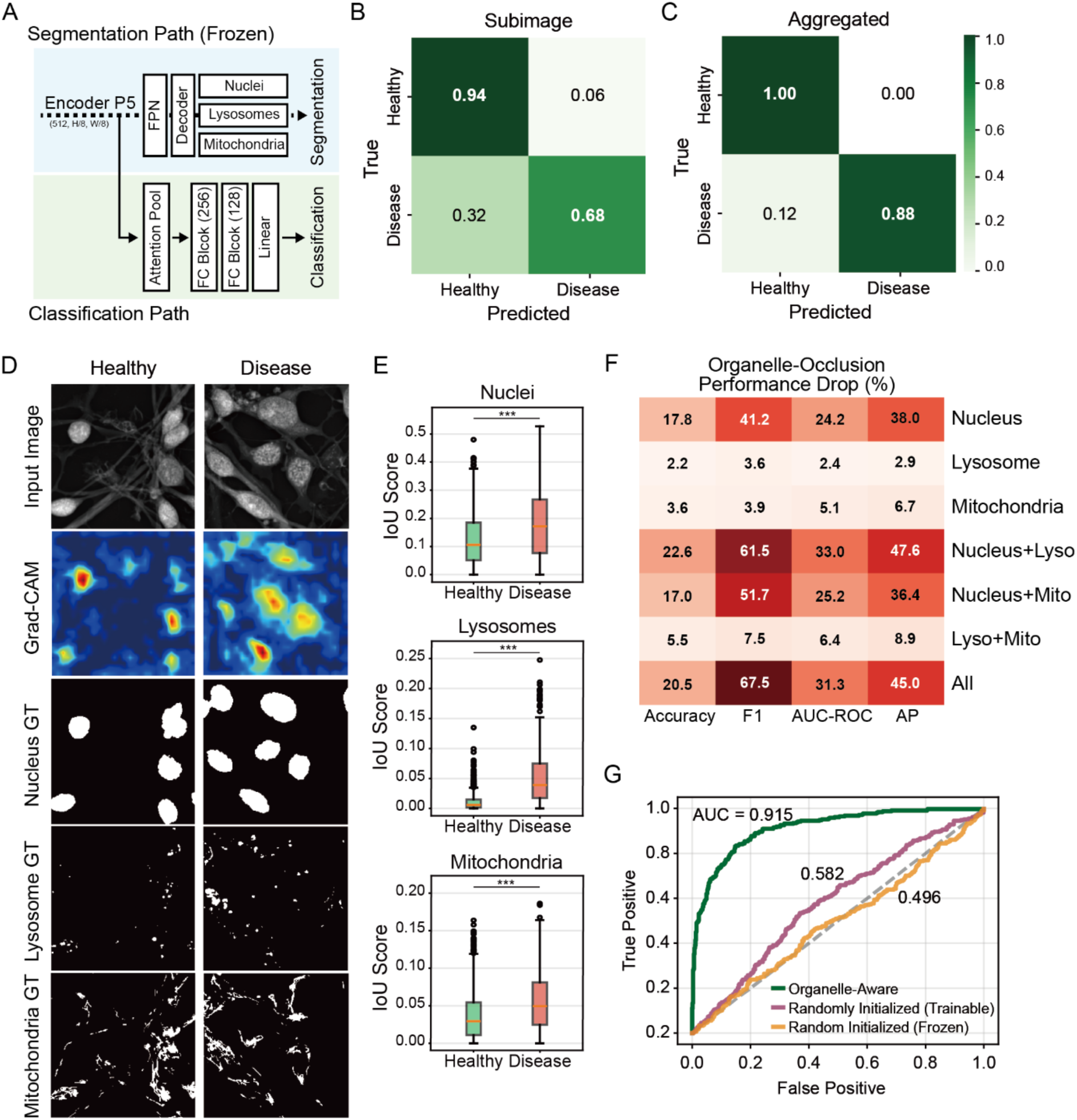
Mitochondrial Dysfunction Classification Leveraging Organelle-Aware Encoder. (A) The classification model architecture overview. The segmentation path is frozen for the classification head training. (B) The end-to-end classification head leveraging a segmentation encoder achieved an overall accuracy of 86% on individual subimages cropped from each full-sized ODT volume. (C) Aggregating subimage classification predictions from the same ODT volume increases the classification accuracy to 96.1%. (D) Grad-CAM examples highlighting the region that contributed towards the model prediction. (E) Correlation quantification of Grad-CAM and organelle ground truths showed that the model focused on each organelle more for the disease-class input than the healthy-class input. This was especially significant for the lysosome. Statistical analysis performed using Mann-Whitney U test with Holm-Bonferroni correction (n=918 subimages, ***: p < 0.001) (F) Organelle-occlusion test demonstrating the impact of masking the organelle region in the input image on classification performance. (G) ROC curves and AUC comparing the three model variants: (i) organelle-aware with pretrained segmentation encoder, (ii) randomly initialized encoder kept frozen, and (iii) fully trainable randomly initialized encoder.

The classifier achieved an accuracy of 85.7% (Fig. 4B), F1 of 0.761 and AUC-ROC (area under receiver operating characteristic curves) of 0.921. During training and evaluation, each full-sized ODT volume was cropped to 7 × 224 × 224 and these metrics were evaluated per crop (subimage). While we adopted intensity(RI)-based filtering at preprocessing step, not all subimage contained valid information in its field of view and subimage classification accuracy does not represent the full ODT volume prediction performance. To overcome this, we aggregated total of 9 subimage predictions (3 × 3) and took the majority voted as the final prediction which resulted in the final prediction accuracy of 96.1% with AUC-ROC of 0.997 (Fig. 4C & Supplementary Fig. 3A).

The interpretation of deep learning model is as important as its performance, and we examined salient regions for the model predictions using Grad-CAM (Fig. 4D). Grad-CAM showed 0.38 Pearson correlation coefficient with the 2D projection of the input volume, implying that the model correctly attended to high RI regions (neurons) instead of the background. Importantly, the intersection-over-union (IoU) between Grad-CAM heatmaps and the fluorescent organelle ground truths revealed that the model attended for the organelles, particularly lysosomes, more strongly in diseased state images than the healthy ones (Fig. 4E). Interpretability was further assessed using occlusion tests, in which specific organelles (or combinations of them) were masked from the input volume to measure their impact on classification performance.

Occluding the nucleus reduced prediction accuracy by 17.8%, lysosomes by 2.2% and mitochondria by 3.6% (Fig. 4F). When all three organelles were occluded, model accuracy dropped by 20.5% and F1 by 67.5%. The effect was even more pronounced under subimage aggregation, resulting in a 30.6% decline in accuracy and a complete (100%) loss of F1 performance. Despite the occluded pixels are only < 5% of the total pixels (Supplementary Fig. 3B), this sharp decline in classification performance indicates that all organelles, especially nuclei accounting for the majority of pixels and cell body, contributed to the accurate prediction of the mitochondrial state of the neurons.

To encourage the classification model to leverage organelle-aware representations, we froze the entire segmentation pathway, including the encoder. However, accurate classification could still be achievable through shortcut learning if the dataset contained spurious class-specific cues such as background illumination, cell density, or preparation artifacts. To test this, we trained an identical architecture without loading the pretrained segmentation weights, thereby starting with a randomly initialized encoder that was kept frozen. Training failed completely and the model produced narrow range of prediction probabilities incapable of identifying mitochondrial dysfunction in neurons (Supplementary Fig. 3C). We further asked whether freezing the randomly initialized encoder had restricted model capacity too aggressively, given the ResNet-34 backbone accounts for the 94% of the trainable parameters in the classification pathway. To test this, we trained the encoder as fully trainable, effectively building a classification model that is equally capable as the organelle-aware model but without the pretrained weights. Even under this condition, the randomly initialized model still failed to achieve accurate classification (Fig 4G & Supplementary Fig. 3D).

Together, these results demonstrate that accurate and interpretable classification of mitochondrial dysfunction from label-free ODT depends critically on organelle-aware representation learning. Furthermore, they indicate that the observed performance is achievable only through pretraining via virtual staining, and no class-specific feature that drive shortcut learning is present in our dataset.

### Morphometric Descriptors from Segmentation Masks Predict Directly Interpretable Classification

To complement the deep-learning classifier with a more direct and transparent approach, we trained a logistic regression model, a classical statistical learning method which provides an interpretable framework by estimating the contribution of each feature to the classification outcome, using morphometric descriptors derived from the predicted organelle-segmentation masks. Connected-component analysis yielded descriptors including organelle area, perimeter, eccentricity, refractive index (RI) distribution, and colocalization with other organelles. Analysis of the descriptors showed that nucleus-lysosome overlap, mitochondria-lysosome overlap, lysosome area, and lysosome count were greatly increased for diseased state neurons (Fig. 5C). These descriptors indicating increase in lysosomal activity for autophagy/mitophagy align well with the biological expectation of dysfunctional mitochondria. Furthermore, the descriptors extracted from the ground truth (fluorescent images) masks and prediction (virtual staining) masks were well aligned supporting the fidelity of the segmentation performance and morphometric descriptors derived from it.

**Figure 5.**
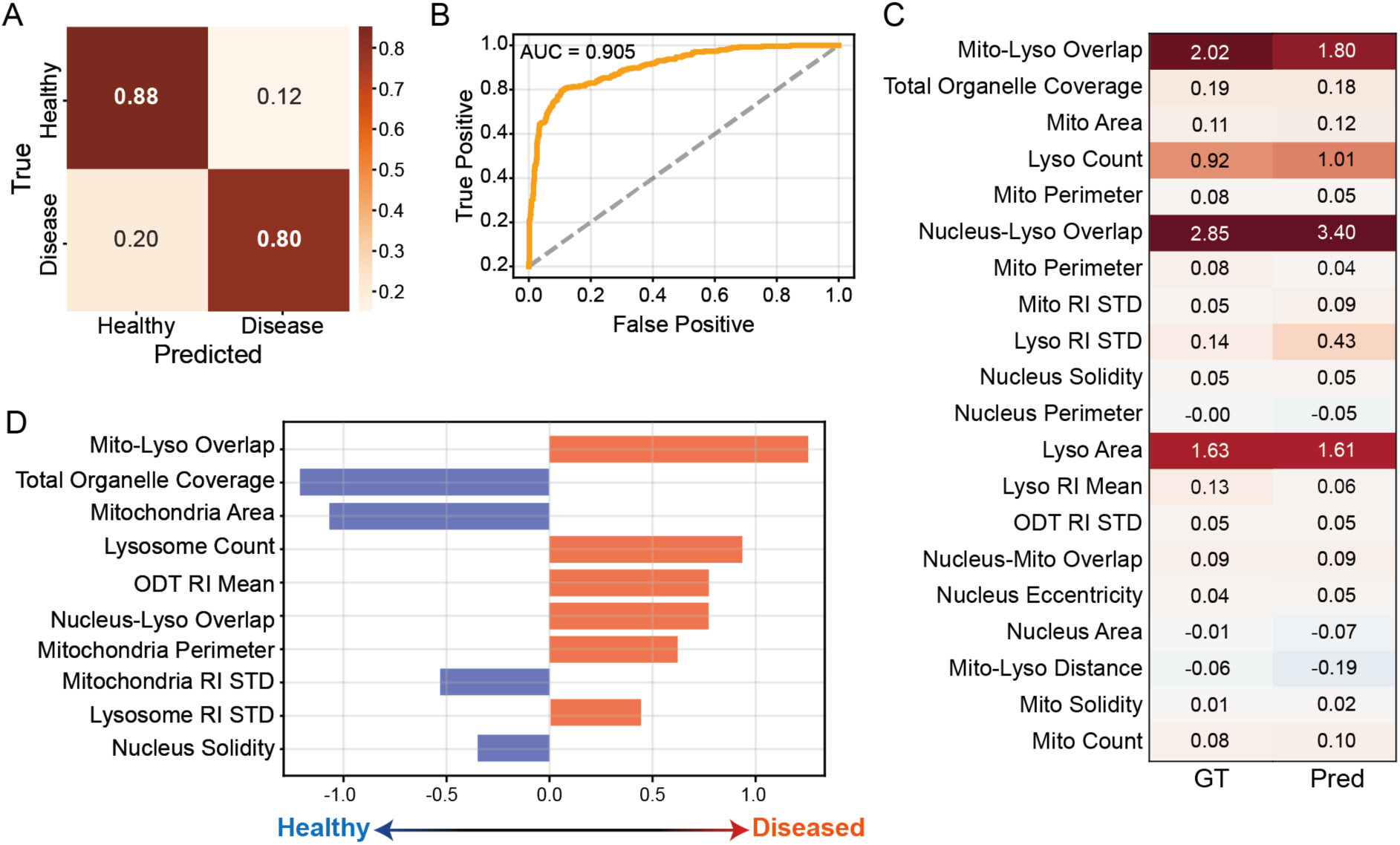
Biologically interpretable classification model trained on morphometric descriptors. (A-B) Logistic regression model trained on morphometric descriptors achieved 82.4% overall accuracy and 0.899 AUC-ROC on the test dataset. (C) The relative difference of morphometric descriptors (disease to healthy). The descriptors extracted using the ground truth masks and the predicted segmentation masks were well aligned, supporting the fidelity of the segmentation prediction-based descriptors (GT: ground truth, Pred: Prediction). (D) Top 10 regression coefficients represent how strongly each feature influences the prediction probability. Positive coefficient features contribute to the diseased state, and negative features contribute to the healthy state. Negative coefficient for mitochondria area indicates a correlation between a small area (fragmentation) and the diseased state.

This model achieved an accuracy of 85.5% and an AUC-ROC of 0.905 (Fig. 5A-B) and 99% accuracy with subimage aggregation (Supplementary Fig. 4A). Analysis of regression coefficients reveals the contribution of each descriptor for the prediction of mitochondrial dysfunction (Fig. 5D & supplementary Fig. 4B). The model identified mitochondria-lysosome overlap as the most informative descriptor (+1.263) which aligns with the biological expectation that increased mitophagy correlates to the mitochondrial dysfunction. Another critical coefficient was mitochondrial area (−1.072). The negative coefficient indicates that smaller mitochondrial cross-section for potential fragmentation strongly favors the prediction of mitochondrial dysfunction. Lysosome count (0.929) and nucleus-lysosome overlap (0.766) were also high-contributors for the prediction which represent increased autophagy.

### Concordance between end-to-end and interpretable models

To evaluate whether our two classification pipelines, (i) the end-to-end classification head by employing the organelle-aware encoder and (ii) the descriptor-based logistic regression model, capture overlapping biological information, we performed a systematic concordance analysis.

Across the identical held-out test set, prediction agreement between the two models was 83.3% and the probability correlation was 0.761, indicating that the interpretable logistic regression model provides meaningful support for the predictions of the relatively black-box deep learning model (Fig. 6A). With subimage aggregation, the probability correlation between the two models increased to 0.945 (Supplementary Fig. 4C). Only 6.5% of the 918 subimages were misclassified by both approaches (Fig. 6B). Direct comparison of each model performance revealed that the accuracy and AUC-ROC were comparable while precision and recall were better for the classification head and logistic regression, respectively (Fig. 6C-D). This suggests that the deep learning method predicts fewer false positives, and the statistical method predicts fewer false negatives. Nevertheless, both approaches support highly accurate and interpretable classification of mitochondrial dysfunction from 3D ODT volume of live hiPSC-derived neurons without any exogeneous labelling.

**Figure 6.**
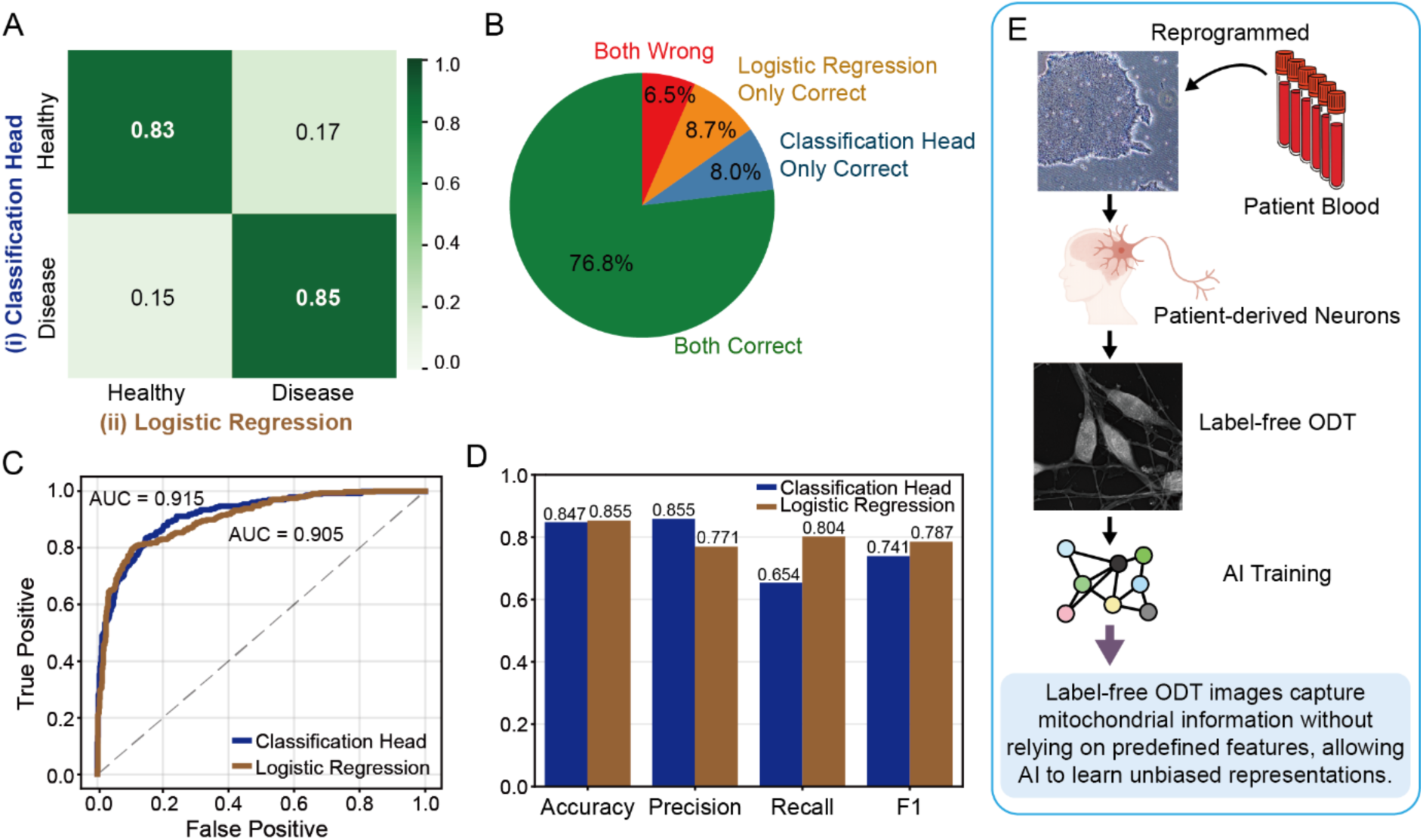
The black-box model aligns with interpretable classification. (A) Prediction agreement between the two classification approaches. This alignment between the end-to-end deep learning approach and the interpretable logistic regression model supports the classification credibility of the deep learning model. (B) Only 6.5% of the 918 test subimages were inaccurately predicted by both models, and 76.8% were classified correctly by both models. (C-D) ROC curve and quantitative metrics comparison showing both the deep learning method and the logistic regression method accurately predict mitochondrial dysfunction from ODT volume. (E) Graphical illustration of the suggested pipeline.

## Discussion

In this study, we demonstrate that optical diffraction tomography (ODT) combined with deep learning can achieve organelle-resolved, label-free analysis of live hiPSC-derived neurons. By implementing a multi-label virtual staining framework for nuclei, mitochondria, and lysosomes, we enabled the encoder to capture organelle-aware representations and extended this capability into two complementary classification pipelines. Together, these approaches accurately and interpretably identify mitochondrial dysfunction in a non-invasive and high-throughput manner.

Interpretability remains a central challenge in applying deep learning to biomedical imaging. We adopt dual-pipeline approach that couples high-capacity, minimal human biased deep learning model and directly interpretable statistical model. The end-to-end head operating on organelle-aware encodings learned via virtual staining achieves robust prediction without engineered features or fluorescent biomarkers that bias the training with human priors. Occlusion and Grad-CAM analyses confirmed that predictions were driven by biologically meaningful regions and ablation analyses showed that the organelle representation learned via virtual staining is critical for the classification performance.

In parallel, the logistic regression model provides direct insights into the mechanistic and morphometric phenotypes. Morphometric descriptors extracted from the predicted organelle masks showed that increased lysosomal activity is observed in diseased neurons. Without any prior knowledge, the simple statistical model identified mitochondria-lysosome engagement (mitochondrial fragmentation, mitophagy, and increased lysosomal activity) as the primary features that distinguish the mitochondrial states. These heuristic findings coincide with the biological expectation of mitochondrial stress.

While this study establishes the feasibility of organelle-resolved virtual staining and classification of mitochondrial dysfunction in live hiPSC-derived neurons, several limitations should be acknowledged. First, virtual staining accuracy is bounded by the fidelity of the ground-truth labels. In our setting, exact voxel-level correspondence between ODT and correlated fluorescence was difficult to guarantee, introducing label noise and spatial misregistration. This effect is most pronounced for lysosomes and mitochondria which are small, sparse, and dynamic targets that occupy only few voxels and are vulnerable to feature loss during early encoder down-sampling. To mitigate this, we augmented the standard ResNet-34 backbone with an additional high-resolution block to preserve fine details but the segmentation performance for these organelles remained modest. Despite this, the downstream classifier was accurate and interpretable, suggesting that organelle-aware encodings distilled from imperfect masks still captured biologically meaningful representations. Second, regarding generalizability, our results demonstrate the organelle representation framework and downstream classification in a single donor cell line. This provides a clear proof of feasibility at the line level and suggests the framework is scalable to additional lines, yet multi-line performance and robustness remain uncertain and will require systematic validation across donors or neuronal subtypes. Finally, although the classification outputs are consistent with early neurodegeneration hallmarks, they are inference-based meaning orthogonal assays must be accompanied to confirm the interpretation fidelity.

By combining deep learning with ODT, we advance label-free imaging toward a quantitative and mechanistic tool for probing mitochondrial health in live neuronal systems. The platform is non-invasive, scalable, and compatible with longitudinal, high-throughput studies, positioning it for immediate use in disease modeling, therapeutic screening, and mechanistic investigations of neurodegeneration. More broadly, the integration of label-free imaging with interpretable representation learning offers a path toward routine, human-minimal profiling of subcellular health in living human neurons, enabling earlier detection of pathology and accelerating the discovery of effective interventions.

## Acknowledgements

This work was supported by the National Research Foundation of Korea, funded by the Korean government (RS-2023-00266872 to M.L.C and D.K, RS-2024-00343012 to M.L.C., RS-2025-00521226 and RS-2025-02223597 to D.K, RS-2024-00440778 and RS-2024-00332454 to K.-J.Y.).

## Data availability

The ODT image dataset has been deposited in Zenodo, with a preview subset available immediately, and the full dataset will be released upon publication of the manuscript. (https://doi.org/10.5281/zenodo.17115497)

## Code availability

The codes used for the models and demonstrations will be available in preview form on GitHub, with the complete repository to be released upon publication (https://github.com/SeungJuYoo-1/ODT-Neuron).

## Methods

Supplementary Table 1.

Supplementary Table 1
|  |  |
| --- | --- |
| Stem Cell Maintenance Medium | mTeSR™ Plus (STEMCELL, 100-0276) |
| Neural Maintenance Medium (N2B27) | DMEM/F-12, Glutamax supplement, 500 ml (Thermo Fisher, 10565018)<br>N2-supplement, 5 ml (Thermo Fisher, 17502048)<br>MEM Non-Essential Amino Acids Solution, 5 ml (Thermo Fisher, 11140050)<br>Penicillin-Streptomycin, 5 ml (Thermo Fisher, 15140122)<br>2-Mercaptoethanol, 1 ml (Sigma-Aldrich, M6250)<br>Insulin solution human, 0.25 ml (Sigma-Aldrich, I9278)<br>Neurobasal medium, minus phenol red, 500 ml (Thermo Fisher, 12348017)<br>B-27 supplement, serum free, 10 ml (Thermo Fisher, 17504044)<br>Glutamax supplement, 5ml (Thermo Fisher, 35050061) |
| Small Molecules and Growth Factors | Y-27632 (Tocris, 1254)<br>bFGF (ThermoFisher, PHG0369)<br>SB431642 (Tocris, 1614)<br>Dorsomorphin dihydrochloride (Tocris, 3093) |
| Antibodies and<br>Fluorescent<br>Probes | Rabbit Anti-MAP2 (Abcam, ab32454)<br>Mouse Anti-SATB1+SATB2 (Abcam, ab51502)<br>Chicken Anti-TUJ1 (GeneTex, GTX85469)<br>Goat Anti-Rabbit, Alexa Fluor 555 (Abcam, ab150078)<br>Goat Anti-Mouse, Alexa Fluor 488 (Abcam, ab150113)<br>Goat Anti-Chicken, Alexa Fluor 647 (Abcam, ab150171) |
| Imaging Dyes | Hoechst 33342 (Thermofisher, 62249)<br>SYTOX Green Nucleic Acid Stain (Invitrogen, S7020)<br>Tetramethylrhodamine, Methyl Ester, Perchlorate (Invitrogen, T668)<br>LysoTracker Deep Red (Invitrogen, L12492)<br>Fluo-4, AM, cell permeant (Invitrogen, F14201) |
| Inducers of<br>Mitochondrial<br>Dysfunctions | Rotenone (Sigma-Aldrich, R8875)<br>Antimycin A from Streptomyces sp. (Sigma-Aldrich, A8674)<br>Oligomycin A (Sigma-Aldrich, 75351) |
| Others | EDTA (Invitrogen, 15575020)<br>Dispase II (Gibco, 17105041)<br>ACCUTASE (STEMCELL, 07920)<br>HBSS, calcium, magnesium, no phenol red (Gibco, 14025092)<br>Triton X-100 (Sigma-Aldrich, X100)<br>BSA (Sigma-Aldrich, A2153)<br>PBS, pH 7.4 (Gibco, 10010015) |

## Stem Cell Maintenance and Neuronal Differentiation

In this study, we used two healthy hiPSC cell lines: SCTi-003A (STEMCELL Technologies), and GM25256 (Coriell Institute). ODT volumes from GM25256 cell line yielded poor quality dataset due to low neuronal differentiation rate and extensive cellular aggregation and were discarded at preprocessing step. All cells were tested negative for mycoplasma (conducted by Cosmogenetech, Daejeon). Stem cells were maintained on a Matrigel (Corning, 354277)-coated plate with StemFlex (Gibco, A3349401) medium. Cells were passaged using 0.5 mM EDTA (Invitrogen, 15575020) when approximately 70% confluent.

The cortical neuron differentiation protocol was adapted from Shi et al. (Fig. 3A). To begin neural induction, stem cells were cultured to over 95% confluency and the StemFlex medium was replaced with N2B27 medium (Supplementary Table 1) supplemented with 10 µM SB431642 (Tocris, 1614) and 1 µM dorsomorphin (Tocris, 3093). The medium with supplements was replaced daily for 10 days. On day 10, cells formed an epithelial sheet and were detached using 1.8 U/ml Dispase (Gibco, 17105041) and split in a 1:2 ratio on Matrigel coated plate with N2B27 medium supplemented with 10 µM Y-27632 (Tocris, 1254) and 1 µM bFGF (Thermo Fisher, PHG0369). The medium was replaced with N2B27 with no additional supplements after 2 days of splitting. Neural rosette appeared around day 15-20. The medium was replaced twice a week, and the cells were passaged every two weeks using Accutase (STEMCELL, 07920) throughout the culture to maintain appropriate cell density and extracellular matrix integrity. Cultured neurons were used in experiments from day 65 to ensure maturity and functionality.

## Immunocytochemistry

Immunocytochemistry (ICC) was performed to confirm the identity and neurodevelopmental properties of the cells. Neurons were fixed with 4% paraformaldehyde for 20 minutes and rinsed three times with phosphate-buffered saline (PBS; Gibco, 10010015). Fixed neurons were permeabilized at room temperature for 1 hour with 0.1% Triton X-100 (Sigma-Aldrich, X100) and 5% bovine serum albumin (BSA; Sigma-Aldrich, A2153) in PBS. Primary antibodies were diluted to a 1:250 working concentration in 5% BSA and incubated with the samples for 1 hour at room temperature or overnight at 4°C. Following three washes with 5% BSA solution, secondary antibodies were applied at a 1:500 working concentration in 5% BSA for 1 hour at room temperature. Nuclei were stained using 10 µM Hoechst 33342 (Thermo Fisher, 62249) in 5% BSA solution for 15 minutes, followed by a single PBS wash. Samples were stored in PBS and imaged with a Nikon AX R confocal microscope.

## Calcium Imaging

The functional properties of the generated neurons were assessed via live-cell calcium imaging using Fluo-4, AM (Invitrogen, F14201) in accordance with the manufacturer’s protocol. Calcium imaging is a widely used technique for evaluating neuronal activity, as intracellular calcium levels serve as a critical secondary messenger in signal transduction pathways and are closely linked to action potential firing and synaptic activity. By forcing calcium responses via membrane depolarization, the functionality of voltage-gated calcium channels can be assessed to validate neuronal health and maturity.

Differentiated cells were seeded at a density of 150,000 cells per well onto an 8-well chamber slide (ibidi, 80827) coated with Matrigel, 24 hours before seeding. Cells were cultured for 7-14 days prior to imaging. The imaging solution was prepared in Hanks’ Balanced Salt Solution (HBSS; Thermo Fisher, 14025092) containing 5 µM Fluo-4. Cells were washed once with HBSS, incubated with the calcium imaging solution for 40 minutes at room temperature, and subsequently washed three times with PBS. Prepared samples were loaded into HBSS and imaged using a Nikon AX R confocal microscope. To induce calcium responses, potassium chloride (KCl; Sigma-Aldrich, P3911) was directly added to the samples at a final concentration of 50 mM.

## Mitochondrial Function Assay

The mitochondrial membrane potential was assessed using Tetramethylrhodamine (TMRM; Invitrogen, T668), a fluorescent dye that accumulates in polarized mitochondria in a membrane potential-dependent manner, providing a reliable indicator of mitochondrial health and functionality. By forcing mitochondrial depolarization, the ability of mitochondria to maintain electrochemical gradients essential for ATP production and calcium homeostasis can be assessed.

Cells were seeded onto 8-well chamber slides following the same preparation protocol as described for calcium imaging. The cells were loaded with HBSS containing 25 nM TMRM for 30 minutes at room temperature. Imaging was performed using a Nikon AX R confocal microscope. A mitochondrial uncoupler, carbonyl cyanide 4-(trifluoromethoxy) phenylhydrazone (FCCP; Sigma-Aldrich, C2920), was added directly to the samples at a final concentration of 10 nM to depolarize mitochondrial membrane potential, and the resultant fluorescent intensity decay was monitored.

## Mitochondrial dysfunction inducing chemical Treatment

Chemical treatments were used to induce specific neuronal dysfunctions to model neurodegenerative disease cellular phenotypes. Each chemical was diluted in appropriate solvent to 1000x the working concentration, and further diluted in N2B27 medium to the desired working concentrations. Cells were washed once with PBS and incubated with the prepared chemical solution for 24 hours at 37°C. Following incubation, cells were washed three times with PBS before proceeding with further experiments.

## Live-cell Organelle Imaging and Quantification

Live-cell imaging was performed to evaluate the effects of chemical treatments on organelle functionality. Four fluorescent dyes were used: Hoechst 33342 (Thermo Fisher, 62249) at 10 µM for nuclei staining, SYTOX Green (Invitrogen, S7020) at 100 nM for permeabilized nuclei, tetramethylrhodamine (Invitrogen, T668) at 25 nM for mitochondria, and LysoTracker Deep Red (Invitrogen, L12492) at 100 nM for lysosomes. These dyes allow for the simultaneous assessment of key organelles and viability markers, enabling comprehensive analysis of cellular responses to treatments.

Cells were prepared on 8-well chamber slides following the same procedure as for calcium imaging. The imaging solution was prepared in HBSS and applied to the cells for 40 minutes at room temperature without washing. Images were captured using a Nikon AX R confocal microscope.

To quantify the area and fluorescent intensity of mitochondria and lysosomes, binary masks were generated for each fluorescent image using adaptive thresholding (opencv-python) to segment and isolate puncta. Colocalization metrics were calculated using Mander’s coefficients, facilitating the evaluation of spatial relationships between mitochondria and lysosomes as an indicator of mitophagy. Cell viability was assessed using Hoechst and SYTOX Green images, with nuclei segmented and counted using the watershed algorithm to distinguish individual cells. One minus the ratio of permeabilized (SYTOX Green-positive) to intact (Hoechst-positive) nuclei served as a quantitative measure of cell viability.

## Optical diffraction tomography acquisition

ODT measurements were acquired using a commercially available tomographic microscope (Tomocube HT-X1, Tomocube Inc., Republic of Korea) as previously described. Differentiated neurons over 65 days were plated onto a 24-well imaging plate (Cellvis, P24-1.5H-N) at 300,000 cells/well. Cells were loaded into HBSS before imaging. For evaluation and analysis purposes, training data were imaged with fluorescent dyes to stain organelles as described for live-cell organelle imaging. Imaging points were manually selected based on cell density within the field of view or randomly selected for filtering at preprocessing step. Obtained raw data were processed for 3D reconstruction and 2D projection using proprietary software (TSX Processing Server, TomoAnalysis) provided by the imaging system manufacturer.

## Segmentation dataset preparation

Segmentation dataset was prepared from all available ODT volumes obtained from other experimental conditions to mitigate data scarcity and improve model generalizability through maximum data availability. This includes chemicals not directly relevant to mitochondrial dysfunction modeling; α-synuclein preformed fibrils (PFF), chloroquine, Imidazole ketone erastin (IKE), RSL3. Segmentation task relies on structural correspondence between ODT volume and corresponding fluorescent labels and is independent phenotype-specific information from these experiments. Only data prepared the same (cell line, culture protocol, imaging configuration and approximate DIV of 60∼80) was pooled to enforce this. Chloroquine (5 µM; Sigma-Aldrich, C6628), IKE (0.5µM; Selleckchem, S8877), and RSL3 (0.5µM; Selleckchem, S8155) was treated using the same method as described for ETC inhibitors for the classification dataset. α-synuclein preformed fibrils (PFF) were kindly provided by the Hyejin Park laboratory (Aging and Neurodegenerative Diseases Laboratory, KAIST). PFF were treated following their previous study^49^ for 3 days at 10 µg/ml and 1 µg/ml.

## Image Preprocessing

Raw ODT volumes were preprocessed through intensity normalization using global minimum and maximum values (13,350-14,000 refractive index units) across the experiments to maintain relative intensity relationships, preserving refractive index information encoded in ODT. Data split was performed per 8:1:1 ratio and the training dataset was augmented including: random rotations (±10°), brightness/contrast adjustments (±8%), Gaussian noise injection (σ=0.015), and minimal geometric distortions. Augmentations were applied consistently to both ODT volumes and corresponding ground truth masks to maintain spatial alignment. Full-sized ODT volumes were cropped as model input preparation to (7, 224, 224). Training dataset was randomly cropped while the validation and test dataset were systematically split into subimages. Finally, input image with extreme low mean intensity values were discarded.

## Segmentation model architecture and training

The segmentation model was implemented as a 2.5D convolutional neural network based on a ResNet-34 encoder with Feature Pyramid Network (FPN) like multi scale fusion layers and organelle-specific decoder architecture. The model processed 7-slice ODT volume inputs (7×224×224) through a 2.5D preprocessing module. The ResNet-34 backbone modified to minimize diminishing small organelle feature was combined with FPN-like multi-scale fusion layers. Three specialized decoder heads generated pixel-wise predictions for each organelle class: (i) Nucleus head with standard convolution layers optimized for large, well-defined nuclear structures; (ii) Lysosome head incorporating multi-scale features and contextual aggregation through dilated convolutions; and (iii) Mitochondria head with similar architecture but optimized for elongated mitochondrial morphology. Each head output class-specific logits processed through sigmoid activation for independent binary classification per pixel. To better handle small and morphologically diverse structures such as lysosomes and mitochondria, an auxiliary heatmap prediction head guides instance separation by learning object centers. The heatmap head predicted Gaussian-smoothed center point heatmaps for each organelle class to improve instance separation quality and contributed to the total loss at 0.5x scale.

Training employed batch size of 8 with mixed-precision computation (AMP), and gradient checkpointing for memory efficiency. The AdamW^50^ optimizer was configured with initial learning rate of 2×10^-4^, weight decay of 1×10^-4^. Learning rate scheduling used cosine annealing with warm restarts (50 epochs) over 500 total epochs until the early stopping triggers based on validation loss improvement.

## Classification model architecture and training

The classification model leveraged the pretrained segmentation encoder while introducing a specialized classification head for binary discrimination between healthy and diseased states. The classification head was appended to the FPN’s highest-level features (P5: 256 channels) and consisted of: (i) global average pooling to aggregate spatial information, (ii) multi-head attention pooling module to capture organelle-specific patterns, and (iii) a multilayer perceptron (512→128→1 neurons) with dropout (p=0.3) and batch normalization for final classification.

All segmentation components (backbone, FPN, decoder, and organelle heads) were frozen with requires_grad=False, ensuring that classification learning relied exclusively on organelle-aware representations while preserving the quality of segmentation features. The training dataset was configured for binary classification by mapping chemical treatments to disease states: healthy controls (Ctrl, n=324) versus mitochondrial dysfunction conditions (Antimycin, Oligomycin, Rotenone, n=696). Data augmentation for classification training included the same structure-preserving transformations used for segmentation, applied only to input ODT volumes. Training configuration follows similar AdamW, learning rate scheduler, AMP, etc.

## Model Interpretability

Model interpretability was assessed using Gradient-weighted Class Activation Mapping (Grad-CAM) computed for the last convolutional layer of the ResNet-34 backbone (layer4[-1].conv2). Occlusion testing evaluated organelle-specific contributions to classification performance. Individual organelle regions were masked using Gaussian blur (σ=3 pixels) while preserving other cellular structures. Classification accuracy degradation quantified the importance of each organelle type for disease discrimination.

## Logistic Regression model

Total of 39 morphometric features were computed across geometric properties and intensity (RI) statistics. These include: area, perimeter, eccentricity, aspect ratio. A regularized logistic regression classifier was trained on the extracted morphometric features using scikit-learn’s LogisticRegression with L2 regularization (C=1.0). Features were standardized using StandardScaler to ensure equal weighting across different measurement scales. The model was trained using the same data splits as the classification head approach.

## Supplementary Materials

**Supplementary Figure 1.**
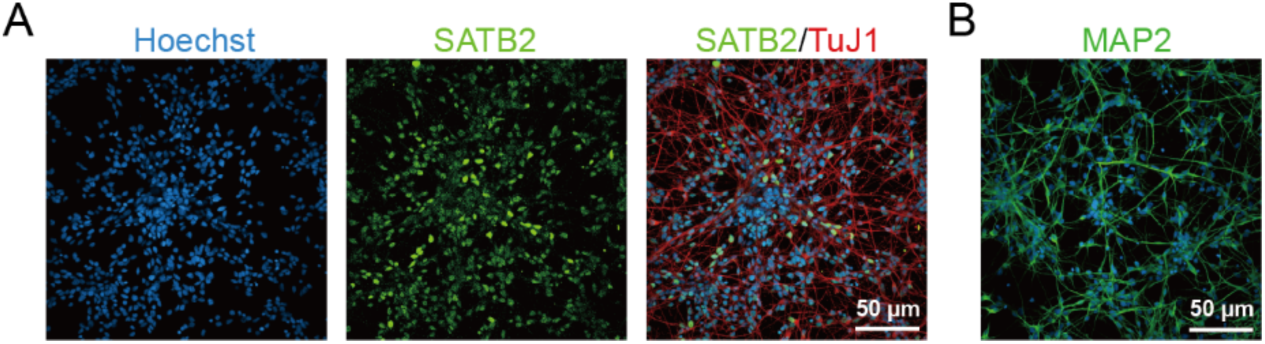
(A-B) Immunocytochemistry of generated hiPSC-derived neurons display strong fluorescence for mature cortical neuron identity.

**Supplementary Figure 2.**
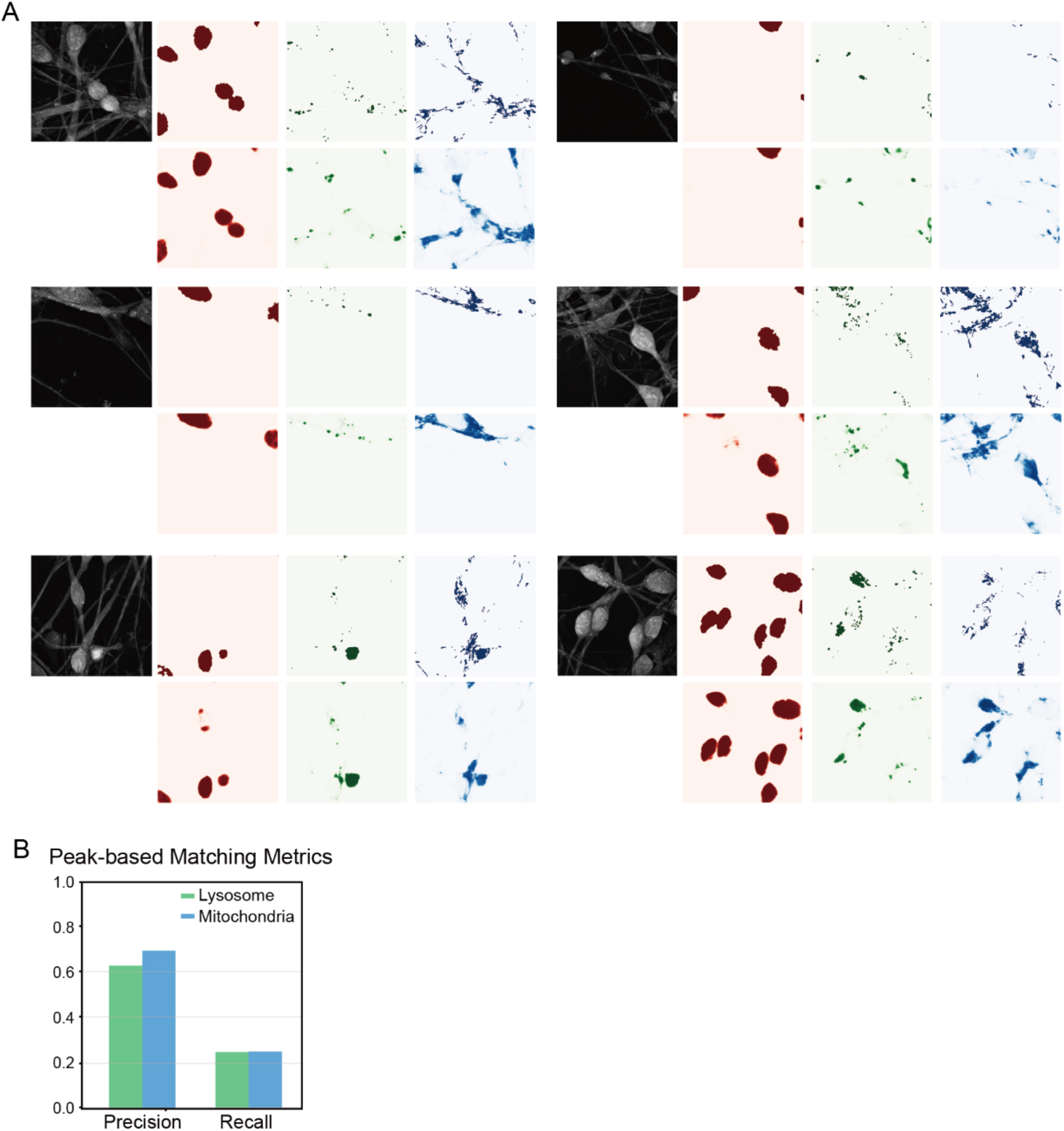
(A) Segmentation examples. Red for nuclei, green for lysosomes, blue for mitochondria. The upper row shows the ground truth from fluorescent imaging, and the lower row shows the predictions. (B) Peak-based instance matching metrics show high precision and low recall for lysosome and mitochondria.

**Supplementary Figure 3.**
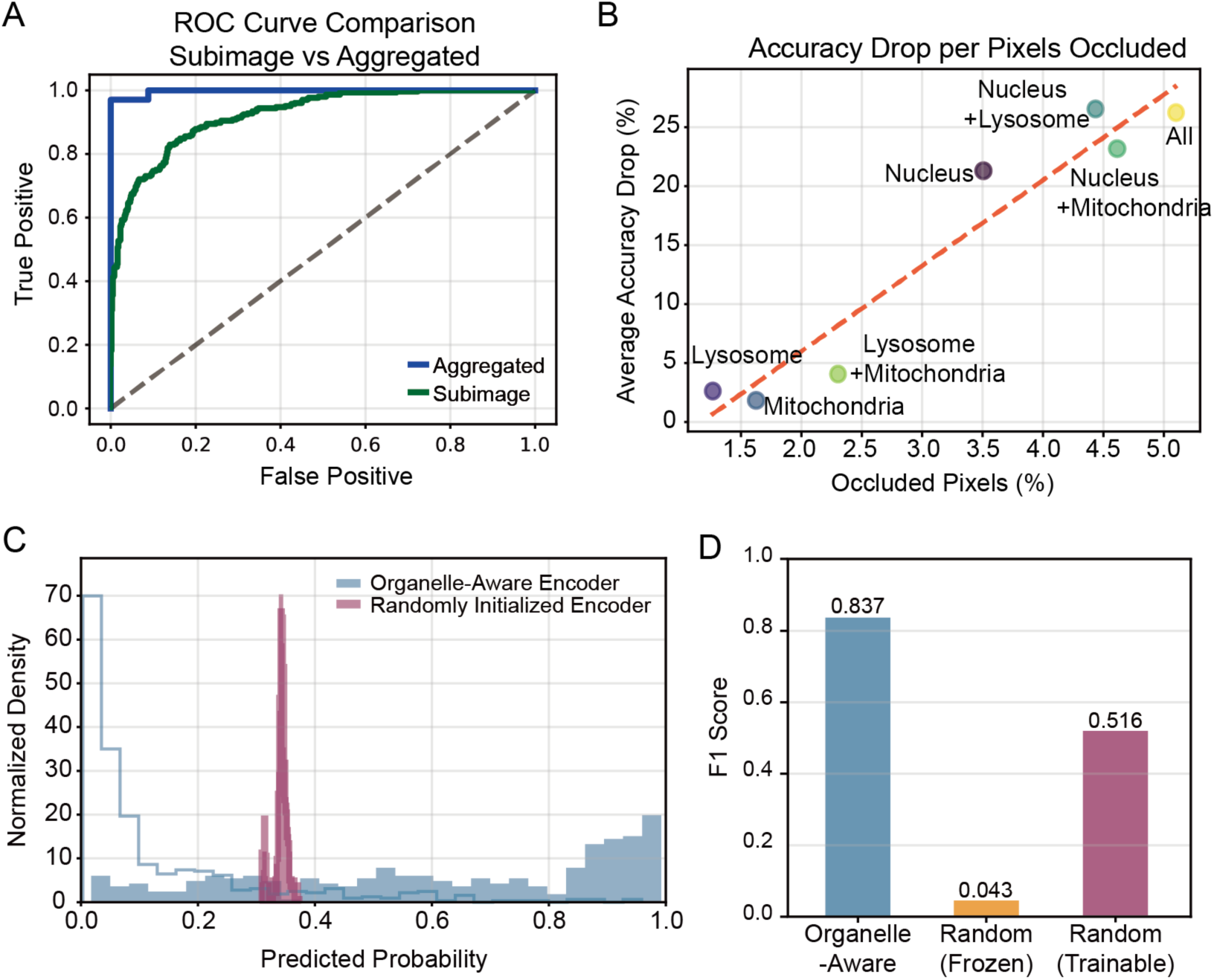
(A) ROC curve comparison for per subimage classification and aggregated classification. During inference, the model takes cropped ODT volume (7 x 224 x 224) for individual subimage prediction. Aggregation utilizes all subimages from the full-size ODT volume for increased robustness and accuracy. (B) Occlusion test show that accuracy drop correlates to the number of pixels occluded. Only 5% occlusion results in 25% decrease in accuracy implying that each organelle contributes significantly to the prediction. (C) Probability distribution comparison between classification models trained using pretrained organelle-aware encoder and randomly initialized encoder. The classification head with randomly initialized encoder instead of pretrained organelle-aware encoder failed to accurately predict mitochondrial dysfunction. (D) F1 score comparison between encoder initialization states after 100 epochs: (i) organelle-aware with pretrained segmentation encoder, (ii) randomly initialized encoder kept frozen, and (iii) randomly initialized encoder trained end-to-end.

**Supplementary Figure 4.**
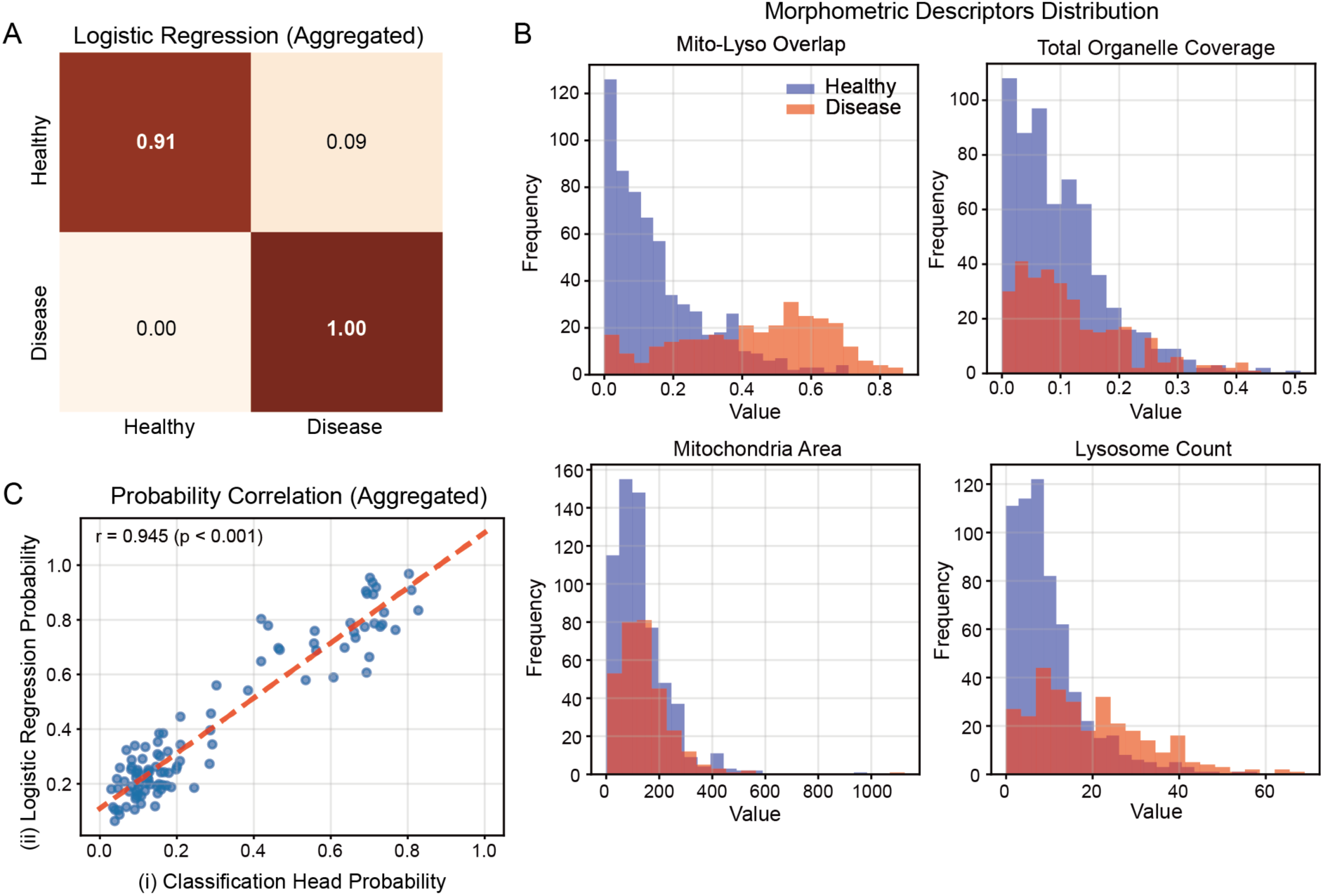
(A) Aggregated classification accuracy for logistic regression model. (B) Top 4 morphometric descriptors distribution. (C) Probability correlation between the two classification approaches. High correlation coefficient supports interpretation of complementary decision making between deep learning method and interpretable logistic regression method.

## References

1. Ben-Shlomo, Y. et al. The epidemiology of Parkinson’s disease. The Lancet 403, 283–292 (2024).

2. Li, X. et al. Global, regional, and national burden of Alzheimer’s disease and other dementias, 1990–2019. Front. Aging Neurosci. 14, (2022).

3. Kim, G. H., Kim, J. E., Rhie, S. J. & Yoon, S. The Role of Oxidative Stress in Neurodegenerative Diseases. Exp. Neurobiol. 24, 325–340 (2015).

4. Lin, M. T. & Beal, M. F. Mitochondrial dysfunction and oxidative stress in neurodegenerative diseases. Nature 443, 787–795 (2006).

5. Cenini, G., Lloret, A. & Cascella, R. Oxidative Stress in Neurodegenerative Diseases: From a Mitochondrial Point of View. Oxid. Med. Cell. Longev. 2019, 2105607 (2019).

6. Kathiresan, D. S. et al. Role of Mitochondrial Dysfunctions in Neurodegenerative Disorders: Advances in Mitochondrial Biology. Mol. Neurobiol. 62, 6827–6855 (2025).

7. Bustamante-Barrientos, F. A. et al. Mitochondrial dysfunction in neurodegenerative disorders: Potential therapeutic application of mitochondrial transfer to central nervous system-residing cells. J. Transl. Med. 21, 613 (2023).

8. D’Alessandro, M. C. B., Kanaan, S., Geller, M., Praticò, D. & Daher, J. P. L. Mitochondrial dysfunction in Alzheimer’s disease. Ageing Res. Rev. 107, 102713 (2025).

9. Singh, T. et al. Neuronal mitochondrial dysfunction in sporadic amyotrophic lateral sclerosis is developmentally regulated. Sci. Rep. 11, 18916 (2021).

10. Wang, W., Zhao, F., Ma, X., Perry, G. & Zhu, X. Mitochondria dysfunction in the pathogenesis of Alzheimer’s disease: recent advances. Mol. Neurodegener. 15, 30 (2020).

11. Toomey, C. E. et al. Mitochondrial dysfunction is a key pathological driver of early stage Parkinson’s. Acta Neuropathol. Commun. 10, 134 (2022).

12. Venkataraman, A. V. et al. Widespread cell stress and mitochondrial dysfunction occur in patients with early Alzheimer’s disease. Sci. Transl. Med. 14, eabk1051 (2022).

13. Takahashi, K. et al. Induction of Pluripotent Stem Cells from Adult Human Fibroblasts by Defined Factors. Cell 131, 861–872 (2007).

14. Kim, K. et al. Epigenetic memory in induced pluripotent stem cells. Nature 467, 285–290 (2010).

15. Park, I.-H. et al. Disease-Specific Induced Pluripotent Stem Cells. Cell 134, 877–886 (2008).

16. Rosenstock, T. R., Sun, C., Hughes, G. W., Winter, K. & Sarkar, S. Analysis of Mitochondrial Dysfunction by Microplate Reader in hiPSC-Derived Neuronal Cell Models of Neurodegenerative Disorders. in Induced Pluripotent Stem Cells and Human Disease: Methods and Protocols (ed. Turksen, K.) 1–21 (Springer US, New York, NY, 2022). doi:10.1007/7651_2021_451.

17. Zatyka, M. et al. Depletion of WFS1 compromises mitochondrial function in hiPSC-derived neuronal models of Wolfram syndrome. Stem Cell Rep. 18, 1090–1106 (2023).

18. Latchman, K., Saporta, M. & Moraes, C. T. Mitochondrial dysfunction characterized in human induced pluripotent stem cell disease models in MELAS syndrome: A brief summary. Mitochondrion 72, 102–105 (2023).

19. Baričević, Z. et al. Label-Free Long-Term Methods for Live Cell Imaging of Neurons: New Opportunities. Biosensors 13, 404 (2023).

20. Ojha, A. & Ojha, N. K. Excitation light-induced phototoxicity during fluorescence imaging. J. Biosci. 46, 78 (2021).

21. Yokoi, Y. et al. Potential consequences of phototoxicity on cell function during live imaging of intestinal organoids. PLOS ONE 19, e0313213 (2024).

22. Kim, G. et al. Holotomography. Nat. Rev. Methods Primer 4, 1–22 (2024).

23. Lim, J. et al. Comparative study of iterative reconstruction algorithms for missing cone problems in optical diffraction tomography. Opt. Express 23, 16933–16948 (2015).

24. Li, J. et al. Optical diffraction tomography microscopy with transport of intensity equation using a light-emitting diode array. Opt. Lasers Eng. 95, 26–34 (2017).

25. Soto, J. M., Rodrigo, J. A. & Alieva, T. Partially Coherent Optical Diffraction Tomography Toward Practical Cell Study. Front. Phys. 9, (2021).

26. Sandoz, P. A., Tremblay, C., Goot, F. G. van der & Frechin, M. Image-based analysis of living mammalian cells using label-free 3D refractive index maps reveals new organelle dynamics and dry mass flux. PLOS Biol. 17, e3000553 (2019).

27. Michael, R., Modirzadeh, T., Issa, T. B. & Jurney, P. Label-Free Visualization and Segmentation of Endothelial Cell Mitochondria Using Holotomographic Microscopy and U-Net. Chem. Biomed. Imaging 3, 225–231 (2025).

28. Chowdhury, S. et al. High-resolution 3D refractive index microscopy of multiple-scattering samples from intensity images. Optica 6, 1211–1219 (2019).

29. Yoon, J. et al. Label-free characterization of white blood cells by measuring 3D refractive index maps. Biomed. Opt. Express 6, 3865–3875 (2015).

30. Pack, C.-G. Application of quantitative cell imaging using label-free optical diffraction tomography. Biophys. Physicobiology 18, 244–253 (2021).

31. Lee, A. J. et al. Label-free monitoring of 3D cortical neuronal growth in vitro using optical diffraction tomography. Biomed. Opt. Express 12, 6928–6939 (2021).

32. Antoine, E. E., Lim, J., Ayoub, A. B., Brandenberg, N. & Psaltis, D. Optical Diffraction Tomography (ODT) for Label-Free Imaging of Large 3D Biological Samples. in Imaging and Applied Optics Congress (2020), paper HF1G.2 HF1G.2 (Optica Publishing Group, 2020). doi:10.1364/DH.2020.HF1G.2.

33. Shi, Y., Kirwan, P. & Livesey, F. J. Directed differentiation of human pluripotent stem cells to cerebral cortex neurons and neural networks. Nat. Protoc. 7, 1836–1846 (2012).

34. D’Sa, K. et al. Prediction of mechanistic subtypes of Parkinson’s using patient-derived stem cell models. *Nat*. Mach. Intell. 5, 933–946 (2023).

35. Li, N. et al. Mitochondrial Complex I Inhibitor Rotenone Induces Apoptosis through Enhancing Mitochondrial Reactive Oxygen Species Production*. J. Biol. Chem. 278, 8516–8525 (2003).

36. Heinz, S. et al. Mechanistic Investigations of the Mitochondrial Complex I Inhibitor Rotenone in the Context of Pharmacological and Safety Evaluation. Sci. Rep. 7, 45465 (2017).

37. Stanford, K. R. et al. Antimycin A-induced mitochondrial dysfunction activates vagal sensory neurons via ROS-dependent activation of TRPA1 and ROS-independent activation of TRPV1. Brain Res. 1715, 94–105 (2019).

38. Stanford, K. R. & Taylor-Clark, T. E. Mitochondrial modulation-induced activation of vagal sensory neuronal subsets by antimycin A, but not CCCP or rotenone, correlates with mitochondrial superoxide production. PLOS ONE 13, e0197106 (2018).

39. Chen, H., Tian, J., Guo, L. & Du, H. Caspase inhibition rescues F1Fo ATP synthase dysfunction-mediated dendritic spine elimination. Sci. Rep. 10, 17589 (2020).

40. Hearne, A., Chen, H., Monarchino, A. & Wiseman, J. S. Oligomycin-induced proton uncoupling. Toxicol. In Vitro 67, 104907 (2020).

41. Alam, M. & Schmidt, W. J. Rotenone destroys dopaminergic neurons and induces parkinsonian symptoms in rats. Behav. Brain Res. 136, 317–324 (2002).

42. Bisbal, M. & Sanchez, M. Neurotoxicity of the pesticide rotenone on neuronal polarization: a mechanistic approach. Neural Regen. Res. 14, 762–766 (2019).

43. Reis, Y. et al. Multi-Parametric Analysis and Modeling of Relationships between Mitochondrial Morphology and Apoptosis. PLOS ONE 7, e28694 (2012).

44. Glancy, B., Kim, Y., Katti, P. & Willingham, T. B. The Functional Impact of Mitochondrial Structure Across Subcellular Scales. Front. Physiol. 11, (2020).

45. Quintana-Cabrera, R. & Scorrano, L. Determinants and outcomes of mitochondrial dynamics. Mol. Cell 83, 857–876 (2023).

46. Chen, W., Zhao, H. & Li, Y. Mitochondrial dynamics in health and disease: mechanisms and potential targets. Signal Transduct. Target. Ther. 8, 333 (2023).

47. He, K., Zhang, X., Ren, S. & Sun, J. Deep Residual Learning for Image Recognition. Preprint at 10.48550/arXiv.1512.03385 (2015).

48. Lin, T.-Y., et al. Feature Pyramid Networks for Object Detection. Preprint at 10.48550/arXiv.1612.03144 (2017).

49. Kam, T.-I. et al. Poly(ADP-ribose) drives pathologic α-synuclein neurodegeneration in Parkinson’s disease. Science 362, eaat8407 (2018).

50. Loshchilov, I. & Hutter, F. Decoupled Weight Decay Regularization. Preprint at 10.48550/arXiv.1711.05101 (2019).

